# RootPainter: Deep Learning Segmentation of Biological Images with Corrective Annotation

**DOI:** 10.1101/2020.04.16.044461

**Authors:** Abraham George Smith, Eusun Han, Jens Petersen, Niels Alvin Faircloth Olsen, Christian Giese, Miriam Athmann, Dorte Bodin Dresbøll, Kristian Thorup-Kristensen

## Abstract

We present RootPainter, a GUI-based software tool for the rapid training of deep neural networks for use in biological image analysis. RootPainter facilitates both fully-automatic and semi-automatic image segmentation. We investigate the effectiveness of RootPainter using three plant image datasets, evaluating its potential for root length extraction from chicory roots in soil, biopore counting and root nodule counting from scanned roots. We also use RootPainter to compare dense annotations to corrective ones which are added during the training based on the weaknesses of the current model.

## Introduction

Plant research is important because we need to find ways to feed a growing population whilst limiting damage to the environment (1). Plant studies often involve the measurement of traits from images, which may be used in phenotyping for genome-wide association studies (2), comparing cultivars for traditional breeding (3) or testing a hypothesis related to plant physiology (4). Plant image analysis has been identified as a bottleneck in plant research (5). A variety of software exists to quantify plant images (6) but is typically limited to a specific type of data or task such as leaf counting (7), pollen counting (8) or root architecture extraction (9).

Convolutional neural networks (CNNs) represent the state-of-the-art in image analysis and are currently the most popular method in computer vision research. They have been found to be effective for tasks both in plant image analysis (10–13) and agricultural research (14, 15).

Developing a CNN-based system for a new image analysis task or dataset is challenging because dataset design, model training and hyper-parameter tuning are time-consuming tasks requiring competencies in both programming and machine learning.

Three questions that need answering when attempting a supervised learning project such as training a CNN are: how to split the data between training, validation and test datasets; how to manually annotate or label the data; and how to decide how much data needs to be collected, labelled and used for training in order to obtain a model with acceptable performance. The choice of optimal hyper-parameters and network architecture are also considered to be a ‘black art’ requiring years of experience and a need has been recognised to make the application of deep learning easier in practice (16).

The question of how much data to use in training and validation is explored in theoretical work that gives indications of a model’s generalisation performance based on dataset size and number of parameters (17). These theoretical insights may be useful for simpler models but provide an inadequate account of the behaviour of CNNs in practice (18).

Manual annotation may be challenging as proprietary tools may be used which are not freely available (19) and can increase the skill set required. Creating dense per-pixel annotations for training is often a time consuming process. It has been argued that tens of thousands of images are required, making small scale plant image datasets unsuitable for training deep learning models (7).

The task of collecting datasets for the effective training of models is further confounded by the unique attributes of each dataset. All data are not created equal, with great variability in the utility of each annotated pixel for the model training process (20). It may be necessary to add harder examples after observing weaknesses in an initial trained model (21), or to correct for a class imbalance in the data where many examples exist of a majority class (22).

Interactive segmentation methods using CNNs such as (23, 24) provide ways to improve the annotation procedure by allowing user input to be used in the inference process and can be an effective way to create large high quality datasets in less time (25).

When used in a semi-automatic setting, such tools will speed up the labelling process but may still be unsuitable for situations where the speed and consistency of a fully automated solution is required. For example when processing data from large scale root phenotyping facilities such as (26) where in the order of 100,000 images or more need to be analysed.

In this study we present and evaluate our software *Root-Painter* which makes the process of creating a dataset, training a neural network and using it for plant image analysis accessible to ordinary computer users by facilitating all required operations with a cross-platform, open-source and freely available user-interface. The RootPainter software was initially developed for quantification of roots in images from rhizotron based root studies. However, we found its versatility to be much broader, with an ability to be trained to recognise many different types of structures in a set of images.

RootPainter allows a user to inspect model performance during the annotation process so they can make a more informed decision about how much and what data is necessary to label in order to train a model to an acceptable accuracy. It allows annotations to be targeted towards areas where the current model shows weakness in order to streamline the process of creating a dataset necessary to achieve a desired level of performance. RootPainter can operate in a semi-automatic way, with a user assigning corrections to each segmented image, whilst the model learns from the assigned corrections, reducing the time-requirements for each image as the process is continued. It can also operate in a fully-automatic way by either using the model generated from the interactive procedure to process a larger dataset without required interaction, or in a more classical way by using a model trained from dense per-pixel annotations which can also be created via the user interface.

We evaluate the effectiveness of RootPainter by training models for three different types of data and tasks without dataset-specific programming or hyper-parameter tuning. We evaluate the effectiveness on a set of rhizotron root images, and in order to evaluate the versatility of the system, also on two other types of data, a biopores dataset, and a legume root nodules dataset, both involving objects in the images quite different from roots.

For each dataset we compare the performance of models trained using the dense and corrective annotation strategies on images not used during the training procedure. If annotation is too time-consuming, then RootPainter will be unfeasible for many projects. To investigate the possibility of rapid and convenient model training we use no prior knowledge and restrict annotation time to a maximum of two hours for each model. We hypothesize that (1) in a limited time period RootPainter will be able to segment the objects of interest to an acceptable accuracy in three datasets including roots, biopores and root nodules, demonstrated by a strong correlation between the measurements obtained from RootPainter and manual methods. And (2) a corrective annotation strategy will result in a more accurate model compared to dense annotations, given the same time for annotation.

Prior work for interactive training for segmentation includes (27) and (28). (27) evaluated their method using neuronal structures captured using Electron Microscopy, and found the interactively trained model to produce better segmentations than a model trained using exhaustive ground truth labels. (28) combined interactive segmentation with interactive training by using the user feedback in model updates. Their training approach requires an initial dataset with full groundtruth segmentations, whereas our method requires no prior labelled data, which was a design choice we made to increase the applicability of our method to plant researchers looking to quantify new objects in a captured image dataset.

As opposed to (27) we use a more modern, fully convolutional network model, which we expect to provide substantial efficiency benefits when dealing with larger images. Our work is novel in that we evaluate an interactive corrective annotation procedure in terms of annotation time to reach a certain accuracy on real-world plant image datasets. Synthetic data is often used to evaluate interactive segmentation methods (29–31). To provide more realistic measurements of annotation time we use real human annotators for our experiments.

### Roots in Soil

Plant roots are responsible for uptake of water and nutrients. This makes understanding root system development critical for the development of resource efficient crop production systems. For this purpose, we need to study roots under real life conditions in the field, studying the effects of crop genotypes and their management (32, 33), cover crops (34), crop rotation (35) and other factors. We need to study deep rooting, as this is critical for the use of agriculturally important resources such as water and nitrogen (36, 37).

Rhizotron based root research is an important example of plant research. Acquisition of root images from rhizotrons is widely adopted (38), as it allows repeated and non-destructive quantification of root growth and often to the full depth of the root systems. Traditionally the method for root quantification in such studies involves a lengthy procedure to determine the root density on acquired images by counting intersections with grid-lines (39).

Manual methods require substantial resources and can introduce undesired inter-annotator variation on root density, therefore a faster and more consistent method is required. More recently, fully automatic approaches using CNNs have been proposed (40); although effective, such methods may be challenging to re-purpose to different datasets for root scientists without the required programming expertise. A method which made the re-training process more accessible and convenient would accelerate the adoption of CNNs within the root research community.

### Biopores

Biopores are tubular or round-shaped continuous voids formed by root penetration and earthworm movement (41). They function as preferential pathways for root growth (42) and are therefore important for plant resource acquisition (43, 44). Investigation of soil biopores is often done by manually drawing on transparent sheets on an excavated soil surface (45). This manual approach is time consuming and precludes a more in-depth analysis of detailed information including diameter, surface area or distribution patterns such as clustering.

### Root Nodules

Growing legumes with nitrogen-fixing capacity reduces the use of fertilizer (46), hence there is an increased demand for legume-involved intercropping (47) and precropping for carry over effects. Roots of legumes form associations with rhizobia, forming nodules on the roots, where the nitrogen fixation occur. Understanding the nodulation process is important to understand this symbiosis and the nitrogen fixation. However, counting nodules from the excavated roots is a cumbersome and time consuming procedure, especially for species with many small nodules such as clovers (*Trifolium spp.*).

## Method

### Software Implementation

RootPainter uses a client-server architecture, allowing users with a typical laptop to utilise a GPU on a more computationally powerful server. The client and server can be used on the same machine if it is equipped with suitable hardware, reducing network IO overhead. Instructions are sent from the client to server using human-readable JSON (JavaScript Object Notation) format. The client-server communication is facilitated entirely with files via a network drive or file synchronisation application. This allows utilisation of existing authentication, authorisation and backup mechanisms whilst removing the need to setup a publicly accessible static IP address. The graphical client is implemented using PyQt5 which binds to the Qt cross-platform widget toolkit. The client installers for Mac, Windows, and Linux are built using the fman build system which bundles all required dependencies. Image data can be provided as JPEG, PNG or TIF and in either colour or grayscale. Image annotations and segmentations are stored as PNG files. Models produced during the training process are stored in the python pickle format and extracted measurements in comma-separated value (CSV) text files.

A folder referred to as the *sync directory* is used to store all datasets, projects and instructions which are shared between the server and client. The server setup (supplementary note 4) requires familiarity with the Linux command line so should be completed by a system administrator. The server setup involves specification of a sync directory, which must then be shared with users. Users will be prompted to input the sync directory relative to their own file system when they open the client application for the first time and it will be automatically stored in their home folder in a file named *root_painter_settings.json* which the user may delete or modify if required.

#### Creating a Dataset

The *Create training dataset* functionality is available as an option when opening the RootPainter client application. It is possible to specify a source image directory, which may be anywhere on the local file system and whether all images from the source directory should be used or a random sample of a specified number of images. It is also possible to specify the target width and height of one or more samples to take from each randomly selected image; this can provide two advantages in terms of training performance. Firstly, RootPainter loads images from disk many times during training which can for larger images (more than 2000 × 2000 pixels) slow down training in proportion to image size and hardware capabilities. Secondly, recent results (48) indicate that capturing pixels from many images is more useful than capturing more pixels from each image when training models for semantic segmentation, thus when working with datasets containing many large images, using only a part of each image will likely improve performance given a restricted time for annotation.

When generating a dataset, each image to be processed is evaluated for whether it should be split into smaller pieces. If an image’s dimensions are close to the target width and height then the image will be added to the dataset without it being split. If an image is substantially bigger then all possible ways to split the image into equally sized pieces above the minimum are evaluated. For each of the possible splits, the resultant piece dimensions are evaluated in terms of their ratio distance from a square and distance from the target width and height. The split which results in the smallest sum of these two distances is then applied. From the split image, up to the *maximum tiles per image* are selected at random and saved to the training dataset. The source images do not need to be the same size and the images in the generated dataset will not necessarily be the same size but all provided images must have a width and height of at least 572 pixels, and we recommend at least 600 as this will allow random crop data augmentation. The dataset is created in the RootPainter sync directory in the datasets folder in a subdirectory which takes the user-specified dataset name. To segment images in the original dimensions, the dataset creation routine can be bypassed by simply copying or moving a directory of images into a subdirectory in the RootPainter datasets directory.

#### Working with Projects

Projects connect datasets with models, annotations, segmentations and messages returned from the server. They are defined by a project file (.seg_proj) which specifies the details in JSON and a project folder containing relevant data. The options to create a project or open an existing project are presented when opening the Root-Painter client application. Creating projects requires specifying a dataset and optionally an initial model file. Alternatively a user may select ‘random weights’ also known as training from scratch, which will use He initialization (49) to assign a models initial weights. A project can be used for inspecting the performance of a model on a given dataset in the client, or training a model with new annotations which can also be created using drawing tools in the client user interface.

#### Model architecture

We modified the network architecture from (40) which is a variant of U-Net (50) implemented in PyTorch (51) using Group Normalization (52) layers. U-Net is composed of a series of down-blocks and up-blocks joined by skip connections. The entire network learns a function which converts the input data into a desired output representation, e.g. from an image of soil to a segmentation or binary map indicating which of the pixels in the image are part of a biopore. In the down-blocks we added 1 × 1 convolution to halve the size of the feature maps. We modified both down-blocks and up-blocks to learn residual mappings, which have been found to ease optimization and improve accuracy in CNNs (53) including U-Net (54). To speed up inference by increasing the size of the output segmentation, we added 1 pixel padding to the convolutions in the down-blocks and modified the input dimensions from 512 × 512 × 3 to 572 × 572 × 3, which resulted in a new respective output size of 500 × 500 × 2, containing a channel for the foreground and background predictions. The modified architecture has approximately 1.3 million trainable parameters, whereas the original had 31 million. These alterations reduced the saved model size from 124.2 MB (55) to 5.3 MB, making it small enough to be conveniently shared via email.

#### Creating Annotations

Annotations can be added by drawing in the user interface with either the foreground (key Q) or background (key W) brush tools. It’s also possible to undo (key Z) or redo brush strokes. Annotation can be removed with the eraser tool (key E). If an image is only partially annotated then only the regions with annotation assigned will be used in the training. Holding the alt key while scrolling can be used to alter the brush size and holding the command key (or windows key) will pan the view. Whilst annotating it’s possible to hide and show the annotation (key A), image (key I) or segmentation (key S). When the user clicks *Save & next* in the interface, the current annotation will be saved and synced with the server, ready for use in training. The first and second annotations are added to the training and validation sets respectively (see *Training Procedure* below). Afterwards, to maintain a typical ratio between training and validation set, annotations will be added to the validation set when the training set is at least five times the size of the validation set, otherwise they will be added to the training set.

#### Training Procedure

The training procedure can be started by selecting *Start training* from the network menu which will send a JSON instruction to the server to start training for the current project. The training will only start if the project has at least two saved annotations as at least one is required for each of the training and validation set. Based on (40) we use a learning rate of 0.01 and Nestorov momentum with a value of 0.99. We removed weight decay as results have shown similar performance can be achieved with augmentation alone whilst reducing the coupling between hyperparameters and dataset (56). The removal of weight decay has also been suggested in practical advice (57) based on earlier results (58) indicating its superfluity when early stopping is used. We do not use a learning rate schedule in order to facilitate an indefinitely expanding dataset.

An *epoch* typically refers to a training iteration over the entire dataset (59). In this context we initially define an epoch to be a training iteration over 612 image sub-regions corresponding to the network input size, which are sampled randomly from the training set images with replacement. We found an iteration over this initial epoch size to take approximately 30 seconds using two RTX 2080 Ti GPUs with an automatically selected batch size of 6. If the training dataset expands beyond 306 images, then the number of sampled sub-regions per epoch is set to twice the number of training images, to avoid validation overwhelming training time. The batch size is automatically selected based on total GPU memory and all GPUs will be used by default using data parallelism.

After each epoch, the model predictions are computed on the validation set and *F*_1_ is calculated for the current and previously saved model. If the current model’s *F*_1_ is higher than the previously saved model then it is saved with its number and current time in the file name. If for 60 epochs no model improvements are observed and no annotations are saved or updated then training will stop automatically.

We designed the training procedure to have minimal RAM requirements which do not increase with dataset size, in order to facilitate training on larger datasets. We found the server application to use less than 8GB of RAM during training and inference, and would suggest at least 16GB RAM for the machine running the server application. We found the client to use less than 1GB RAM but have not yet tested on devices equipped with less than 8GB of RAM.

#### Augmentation

We modified the augmentation procedure from (40) in three ways. We changed the order of the transforms from fixed to random in order to increase variation. We reduced the probability that each transform is applied to 80% in order to reduce the gap between clean and augmented data, which recent results indicate can decrease generalization performance (60). We also modified the elastic grid augmentation as we found the creation of the deformation maps to be a performance bottleneck. To eliminate this bottleneck we created the deformation maps at an eighth of the image size and then interpolated them up to the correct size.

#### Creating Segmentations

It is possible to view segmentations for each individual image in a dataset by creating an associated project and specifying a suitable model. The segmentations are generated automatically via an instruction sent to the server when viewing each image and saved in the segmentations folder in the corresponding project.

When the server generates a segmentation, it first segments the original image and then a horizontally flipped version. The output segmentation is computed by taking the average of both and then thresholding at 0.5. This technique is a type of test time data augmentation which is known to improve performance (61). The segmentation procedure involves first splitting the images into tiles with a width and height of 572 pixels, which are each passed through the network and then an output corresponding to the original image is reconstructed.

It’s possible to segment a larger folder of images using the *Segment folder* option available in the network menu. To do this, an input directory, output directory and one or more models must be specified. The model with the highest number for any given project will have the highest accuracy in terms of *F*_1_ on the automatically selected validation set. Selecting more than one model will result in model averaging, an ensemble method which improves accuracy as different models don’t usually make identical errors (59). Selecting models from different projects representing different training runs on the same dataset will likely lead to a more diverse and thus more accurate ensemble, given they are of similar accuracy. It it is also possible to use models saved at various points from a single training run, a method which can provide accuracy improvements without extending training time (62).

#### Extracting Measurements

It is possible to extract measurements from the produced segmentations by selecting an option from the measurements menu. The *Extract length* option extracts centerlines using the skeletonize method from scikit-image (63) and then counts the centerline pixels for each image. The *Extract region properties* uses the scikit-image regionprops method to extract the coordinates, diameter, area, perimeter and eccentricity for each detected region and stores this along with the associated filename. The *Extract count* method gives the count of all regions per image. Each of the options require the specification of an input segmentation folder and an output CSV.

### A. Datasets

#### A.1. Biopore Images

Biopore images were collected near Meckenheim (50°37’9″N 6°59’29″E) at a field trial of University of Bonn in 2012 (see (45) for a detailed description). Within each plot an area was excavated to a depth of 0.45 m. The exposed soil surface was carefully flattened to reveal biopores and then photographed.

Bersoft software (Windows, Version 7.25) was used for biopore quantification. Using the eclipse function, the visible biopores were marked, then the count number was generated as a CSV file. Pores smaller than 2 mm were excluded from biopore counting.

We restricted the analysis to images with a suitable resolution and cropped to omit border areas. For each image, the number of pixels per mm was recorded using Gimp (MacOS, Version 2.10) in order to calculate pore diameter. We split the images into two folders. BP_counted which contained 39 images and was used for model validation after training as these images had been counted by a biopore expert and BP_uncounted which contained 54 images and was used for training.

#### A.2. Nodule Images

Root images of persian clover (*Trifolium resupinatum*) were acquired at 800 DPI using a waterbed scanner (Epson V700) after root extraction. We used a total of 113 images which all had a blue background, but were taken with two different lighting settings. From the 113 images, 65 appeared darker and underexposed where as 48 were well lit and appeared to show the nodules more clearly. They were counted manually using WinRhizo Pro (Regent Instruments Inc., Canada, Version 2016). Image sections were enlarged and nodules were selected manually by clicking. Then, the total number of marked nodules were counted by the software. We manually cropped to remove the borders of the scanner using Preview (MacOS, Version 10.0) and converted to JPEG to ease storage and sharing. Of these 50 were selected at random to have subregions included in training and the remaining 63 were used for validation.

#### A.3. Roots Dataset

We downloaded the 867 grid counted images and manual root length measurements from (67) which were made available as part of the evaluation of U-Net for segmenting roots in soil (40) and originally captured as part of a study on chicory drought stress (4) using a 4 m rhizobox laboratory described in (68). We removed the 10 test images from the grid counted images, leaving 857 images. The manual root length measurements are a root intensity measurement per-image, which was obtained by counting root intersections with a grid as part of (4).

### B. Annotation and Training

For the roots, nodules and biopores we created training datasets using the *Create training dataset* option. We used random sample, with the details specified in Table 1. The two users (user a and user b) that we used to test the software were the two first authors. Each user trained two models for each dataset. For each model, the user had two hours (with a 30 minutes break between them) to annotate 200 images. We first trained a model using the corrective annotation strategy whilst recording the finish time and then repeated the process with the dense annotation strategy, using the recorded time from the corrective training as a time limit. This was done to ensure the same annotation time was used for both annotation strategies. With corrective annotations, the annotation and training processes are coupled as there is a feedback loop between the user and model being trained that happens in real time. Whereas with dense the user annotated continuously, without regard to model performance. The protocol followed when using corrective annotations is outlined in Supplementary Note 1 and annotation advice given in Supplementary Note 2. For the first six annotations on each dataset, we added clear examples rather than corrections. This was because we observed divergence in the training process when using corrective from the start in preliminary experiments. We suspect the divergence was caused by the user adding too many background classes compared to foreground or difficult examples. When creating dense annotations, we followed the procedure described in Supplementary Note 3.

**Table 1.** Details for each of the datasets created for training. The number of images and tiles were chosen to enable a consistent dataset size of 200 images. Only 50 images were sampled from for the biopores and nodules, in order to ensure there were enough images left in the test set. The datasets created are available to download from (64).

| Object | Name | Source folder | Source URL | To sample | Max tiles | Target size |
| --- | --- | --- | --- | --- | --- | --- |
| Biopores | BP_750_training | BP_uncounted | (65) | 50 | 4 | 750 |
| Nodules | nodules_750_training | counted_nodules | (66) | 50 | 4 | 750 |
| Roots | towers_750_training | grid_counted_roots | (67) | 200 | 1 | 750 |

When annotating roots, in the interests of efficiency, a small amount of soil covering the root would still be considered as root if it was very clear that root was still beneath. Larger gaps were not labelled as root. Occluded parts of nodules were still labelled as foreground (Figure 2). Only the centre part of a nodule was annotated, leaving the edge as undefined. This was to avoid nodules which were close together being joined into a single nodule. When annotating nodules which were touching, a green line (background labels) was drawn along the boundary to teach the network to separate them so that the segmentation would give the correct counts (Figure 3).

**Fig. 1.**
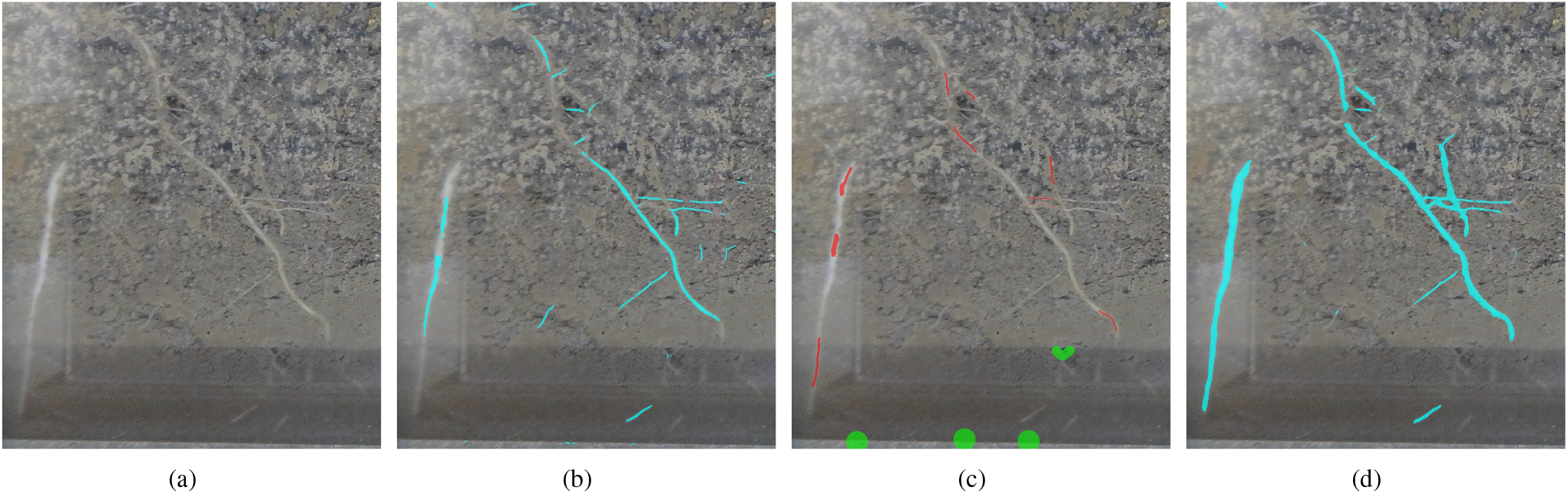
RootPainter corrective annotation concept. (a) Roots in soil. (b) AI root predictions. (c) Human corrections. (d) AI learns from corrections.

**Fig. 2.**
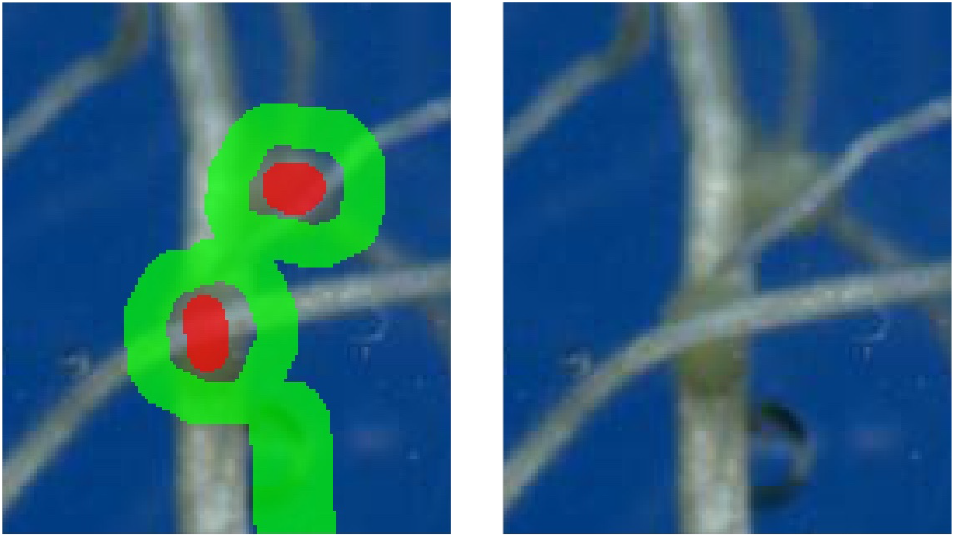
We annotated nodules occluded by roots as though the roots were not there. The red brush was used to mark the foreground (nodules) and the green brush to mark the background (not nodules).

**Fig. 3.**
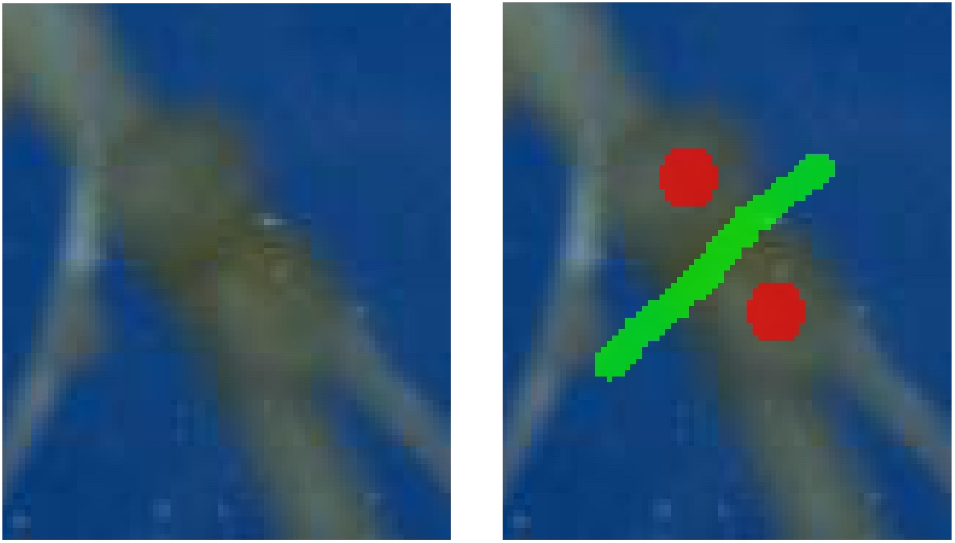
Adjacent nodules were separated using the background class. The red brush was used to mark the foreground (nodules) and the green brush to mark the background (not nodules).

After completing the annotation, we left the models to finish training using the early stopping procedure and then used the final model to segment the respective datasets and produce the appropriate measurements.

We also repeated this procedure for the projects but using a restricted number of annotations by limiting to those that had been created in just 30, 60, 90, 120 and 150 minutes (including the 30 minute break period) to give us an indication of model progression over time with the two different annotation strategies.

### C. Measurement and Correlation

For each project we obtained correlations with manual measurements using the portion of the data not used during training to give a measure of generalization error, which is the expected value of the error on new input (59). For the roots dataset, the manual measurements were compared to length estimates given by RootPainter, which are obtained from the segmentations using skeletonization and then pixel counting.

For the biopores and nodules datasets we used the extract region properties functionality from RootPainter, which gives information on each connected region in an output segmentation. For the biopores the regions less than 2mm in diameter were excluded. The number of connected regions for each image were then compared to the manual counts.

## Results

We report the *R*^2^ for each annotation strategy for each user and dataset (Table 2). Training with corrective annotations resulted in strong correlation (*R*^2^ ≥ 0.7) between the automated measurements and manual measurements five out of six times. The exception was the nodules dataset for user b with an *R*^2^ of 0.69 (Table 2). Training with dense annotations resulted in strong correlation three out of six times, with the lowest *R*^2^ being 0.55 also given by the nodules dataset for user b (Table 2).

**Table 2.** *R*^2^ for each training run. These are computed by obtaining measurements from the segmentations from the final trained model and then correlating with manual measurements for the associated dataset.

| Dataset | User | Corrective $R^2$ | Dense $R^2$ |
| --- | --- | --- | --- |
| Biopores | a | 0.78 | 0.58 |
| Biopores | b | 0.78 | 0.67 |
| Nodules | a | 0.73 | 0.89 |
| Nodules | b | 0.69 | 0.55 |
| Roots | a | 0.89 | 0.90 |
| Roots | b | 0.92 | 0.90 |

For each annotation strategy, we report both the mean and standard error for the obtained *R*^2^ values from all datasets and both users (Table 3). The mean of the *R*^2^ values obtained when using corrective annotation shows they tended to be higher compared with dense, but the differences were not statistically significant (Mixed-effects model; P≤0.05). We plot the mean and standard error at each time point for which multiple *R*^2^ values were obtained (Figure 4). In general corrective improved over time, overtaking dense performance just after the break in annotation (Figure 4). The 30 minute break period taken by the annotator after one hour corresponds to a flat line in performance during that period (Figure 4). On average, dense annotations were more effective at the 30 minute time period, whereas corrective were more effective after two hours (including the 30 minute break) and at the end of the training (Table 3).

**Table 3.** Mean and standard error of the *R*^2^ for each annotation strategy. These are computed by obtaining measurements from the segmentations from the final trained model and then correlating with manual measurements. Using mixed-effects model with annotation strategy as a fixed factor and user and dataset as random factors no significant effects were found (*P* ≤ 0.05).

| Strategy | Mean | Standard error |
| --- | --- | --- |
| Corrective | 0.80 | 0.04 |
| Dense | 0.75 | 0.07 |

**Fig. 4.**
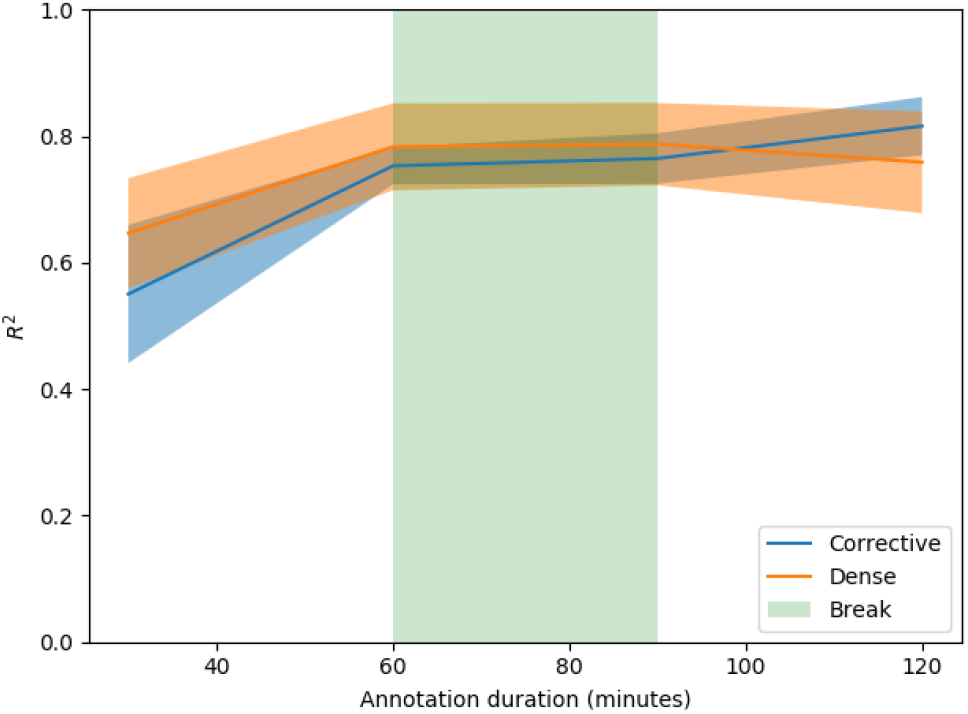
Mean and standard error for the *R*^2^ values over time. These include the 30 minute break and are restricted to time points where multiple observations are available.

We report the duration for each user and dataset (Table 5). Five out of six times all 200 images were annotated in less than the two hour time limit. The nodules dataset took the least time, with annotation completed in 66 minutes and 80 minutes for users a and b respectively (Figure 5). The roots dataset for user a was the only project where the two hours time limit was reached without running out of images (Figure 5).

**Fig. 5.**
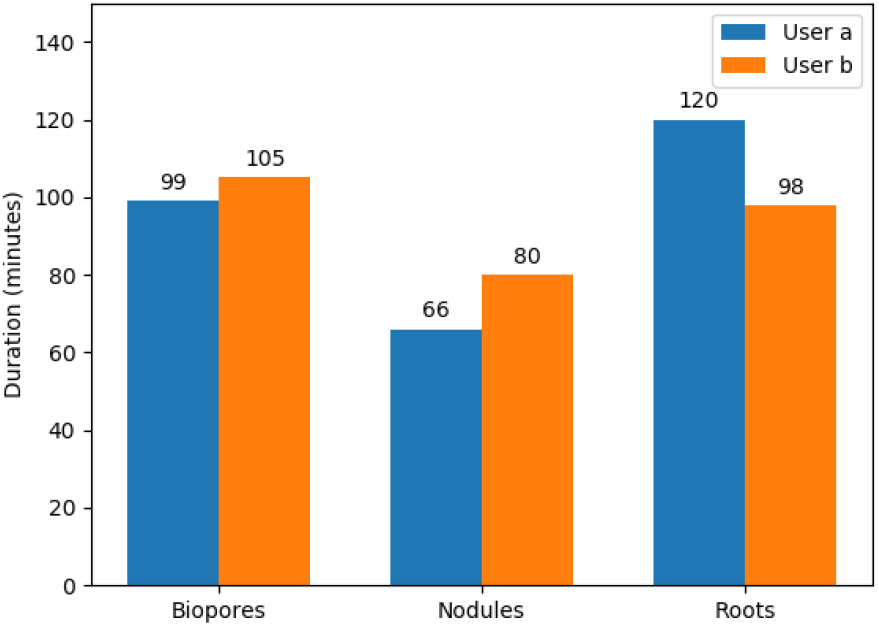
User reported duration in minutes for annotating each dataset, excluding the 30 minutes break taken after one hour of annotation. The annotator would use the same amount of time for both corrective and dense annotation strategies. It fell below the limit of 2 hours (excluding break) when they ran out of images to annotate.

We show an example of errors found from the only model trained correctively which did not result in a strong correlation (Figure 6). There were cases when the vast majority of pixels were labelled correctly but a few small incorrect pixels could lead to substantial errors in count (Figure 6). We show examples of accurate segmentation results obtained with models trained using the corrective annotation strategy (Figures 7, 8 and 9) along with the corresponding manual measurements plotted against the automatic measurements obtained using RootPainter (Figure 10).

**Fig. 6.**
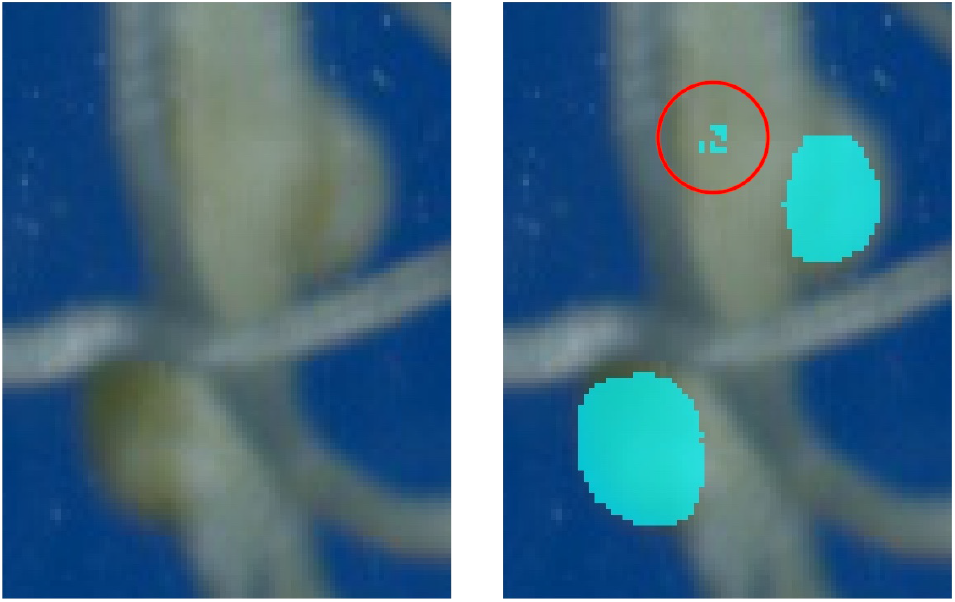
Two correctly detected nodules shown with three false positives. Segmentation is shown overlaid on top of a sub-region of one of the nodule images used for evaluation. The correct nodules are much larger and on the edge of the root. The three false positives are indicated by a red circle. They are much smaller and bunched together.

**Fig. 7.**
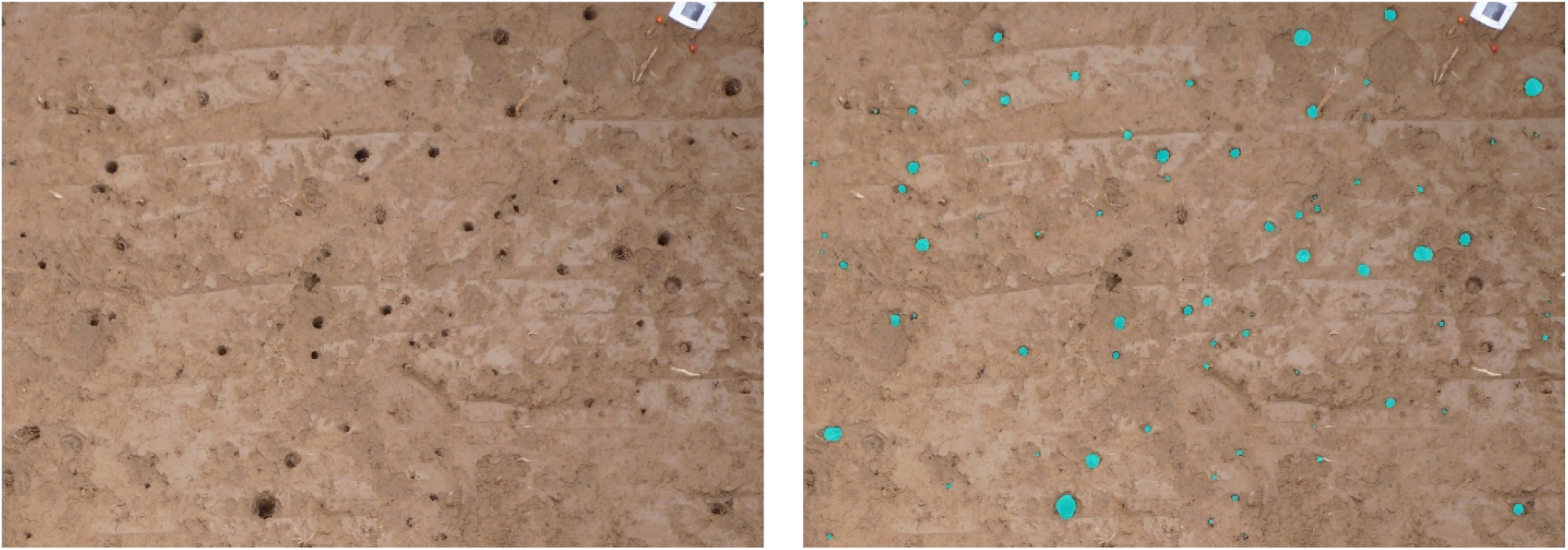
Example input and segmentation output from biopore photographs not used in training. The segmentation was generated using the model trained with corrective annotations by user b. The model was trained from scratch using no prior knowledge with annotations created using RootPainter in 1 hour and 45 minutes.

**Fig. 8.**
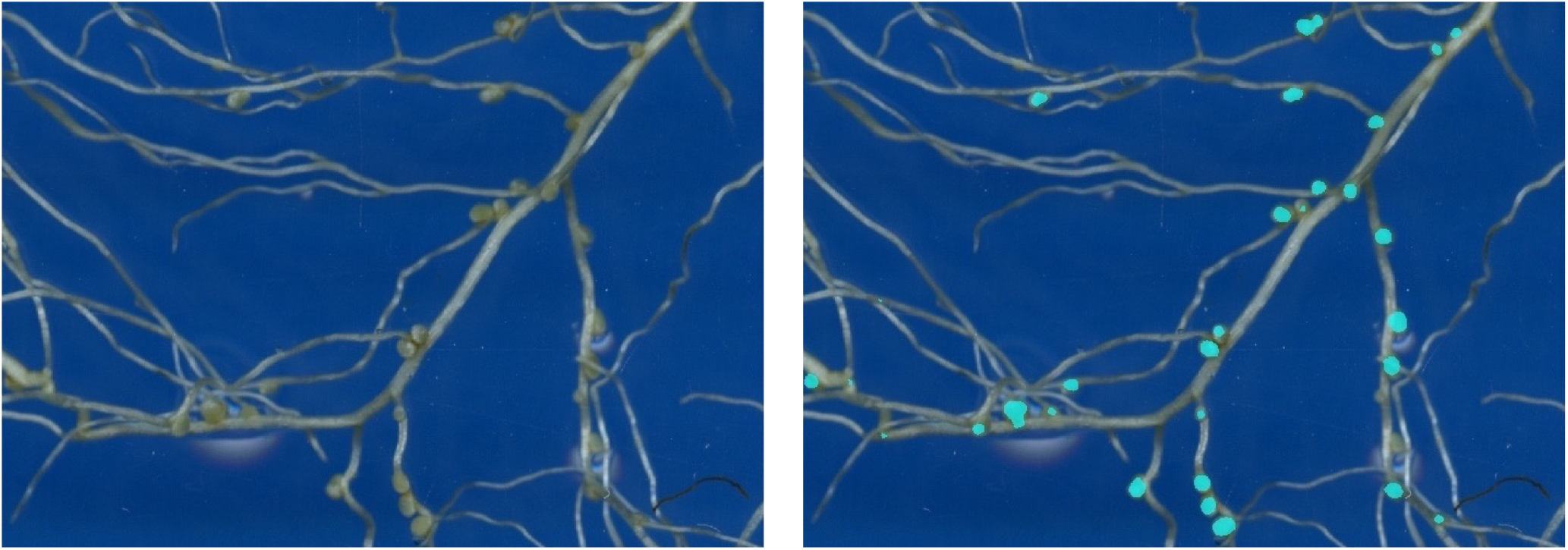
Example input and segmentation output from the nodule scans not used in training. The segmentation was generated using the model trained with corrective annotations by user a. The model was trained from scratch using no prior knowledge with annotations created using RootPainter in 1 hour and 6 minutes.

**Fig. 9.**
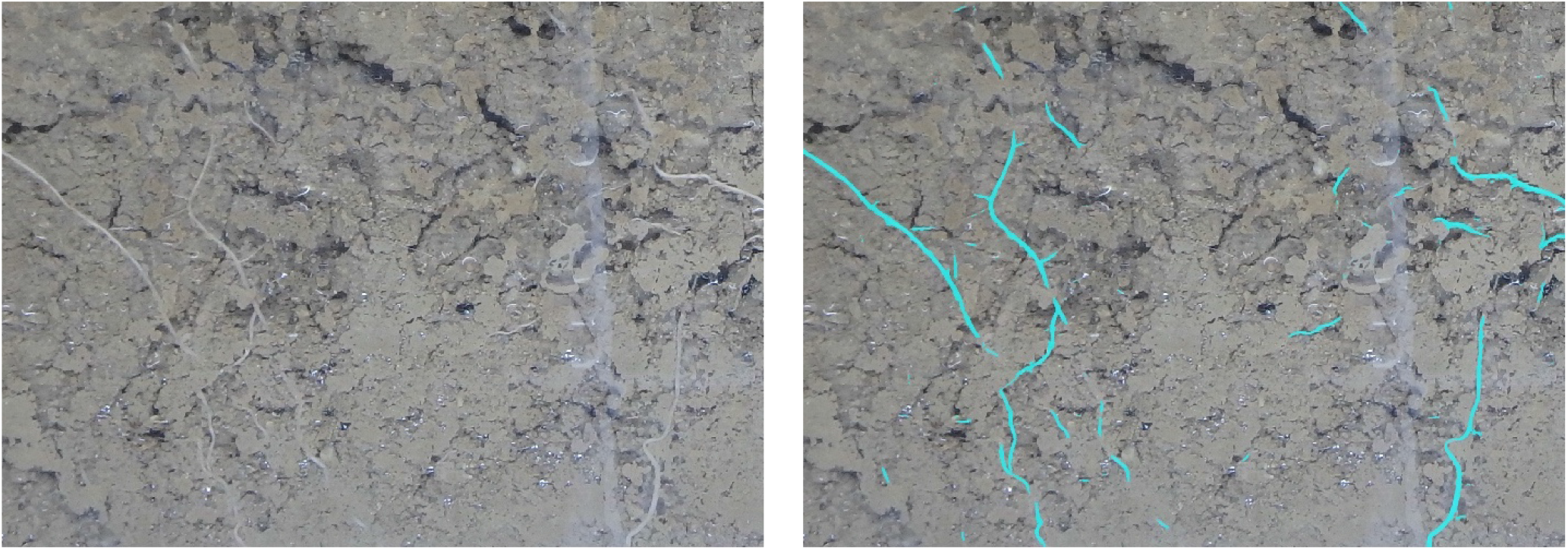
Example input and segmentation output from the grid-counted roots not used in training. The segmentation was generated using the model trained with corrective annotations by user a. The model was trained from scratch using no prior knowledge with annotations created using RootPainter in two hours.

**Fig. 10.**
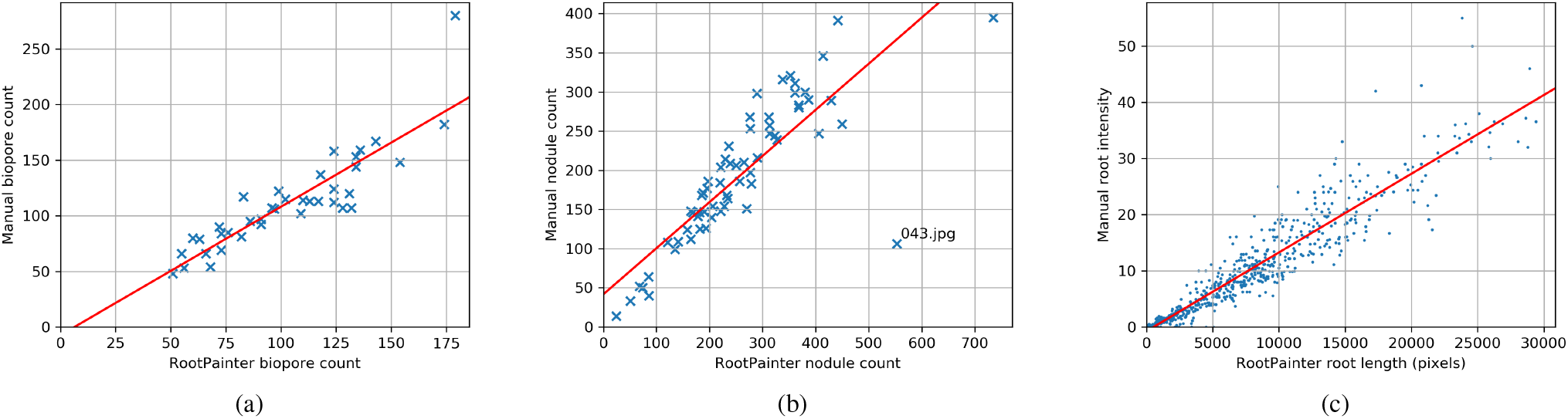
Manual measurements plotted against automatic measurements attained using RootPainter. (a) Biopores using user b corrective model. (b) Nodules using user a corrective model. (c) Roots in soil using user a corrective model.

The observed *R*^2^ values for corrective annotation had a significant positive correlation with annotation duration (P<0.001). There was no significant correlation between annotation time and *R*^2^ values for models trained using dense annotations.

We plot the *R*^2^ for each project after training was completed along with the *R*^2^ obtained with training done only on annotations at restricted time limits, and refer to these as *trained to completion* along with the models saved at that time point during the corrective annotation procedure as it happened which we refer to as *real time* (Figure 11). After only 60 minutes of annotation, all models trained for roots in soil gave a strong correlation with grid counts (Figure 11, Roots a and b). The performance of dense annotation for user b on the nodules dataset was anomalous with a decrease in *R*^2^ as more annotated data was used in training (Figure 11, Nodules b). The corrective models obtained in real time were similar to those trained to completion, except nodules by user b, indicating that computing power was sufficient for real time corrective training (Figure 11).

**Fig. 11.**
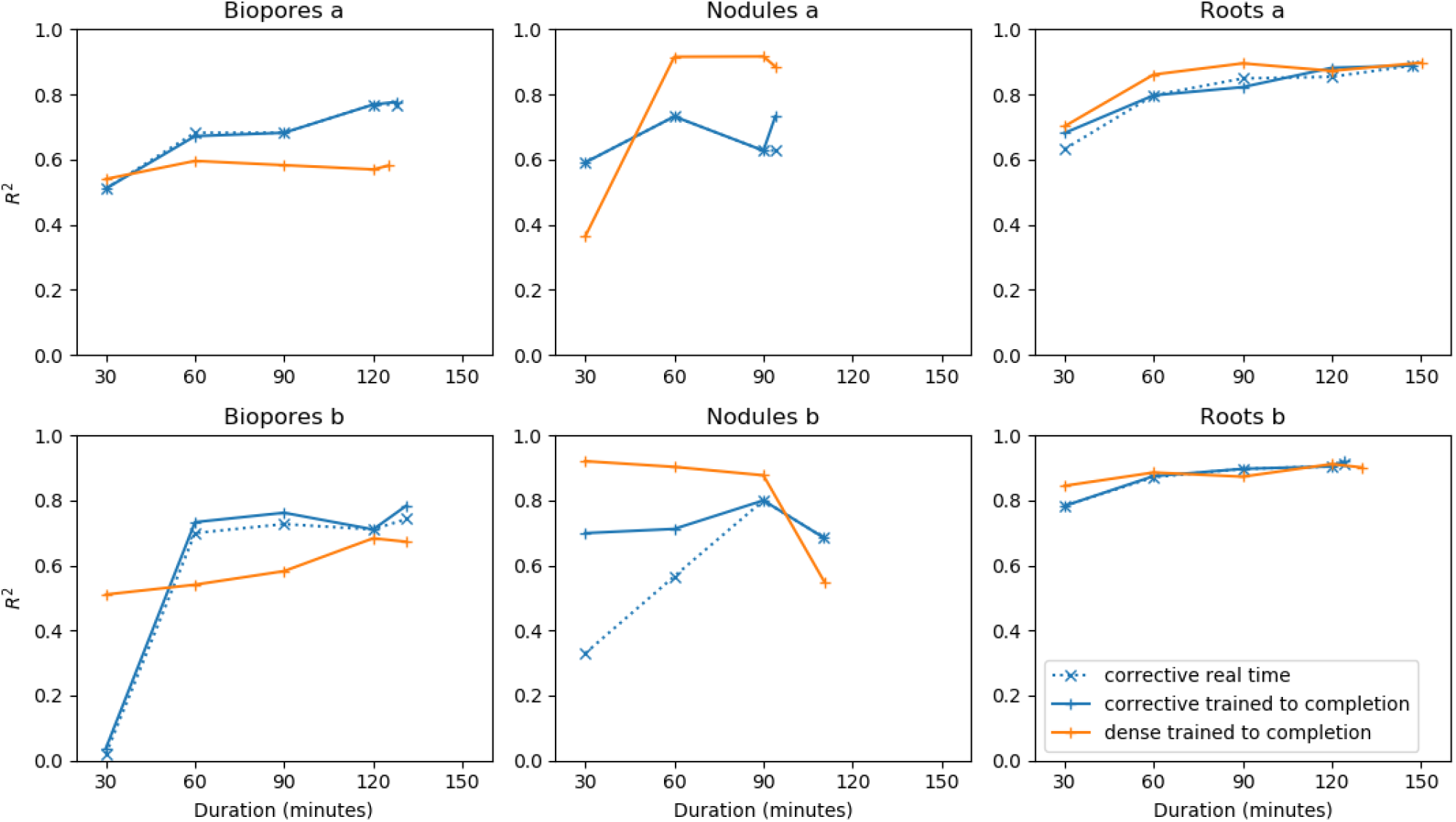
*R*^2^ for the annotations attained after 30, 60, 90, 120 minutes and the final time point for users a and b on the three datasets for dense and corrective annotation strategies. *trained to completion* refers to models which were trained until stopping without interaction, using the annotations created within the specified time period, whereas *real time* refers to models saved during the corrective annotation procedure as it happened. For the corrective annotations we plot both the performance of the model saved during the training procedure and the same model if allowed to train to completion with the annotations available at that time.

We plot the number of images viewed and annotated for the corrective and dense annotation strategies (Figure 12). For the corrective annotation strategy, only some of the viewed images required annotation. In all cases the annotator was able to progress through more images using corrective annotation (Figure 12).

**Fig. 12.**
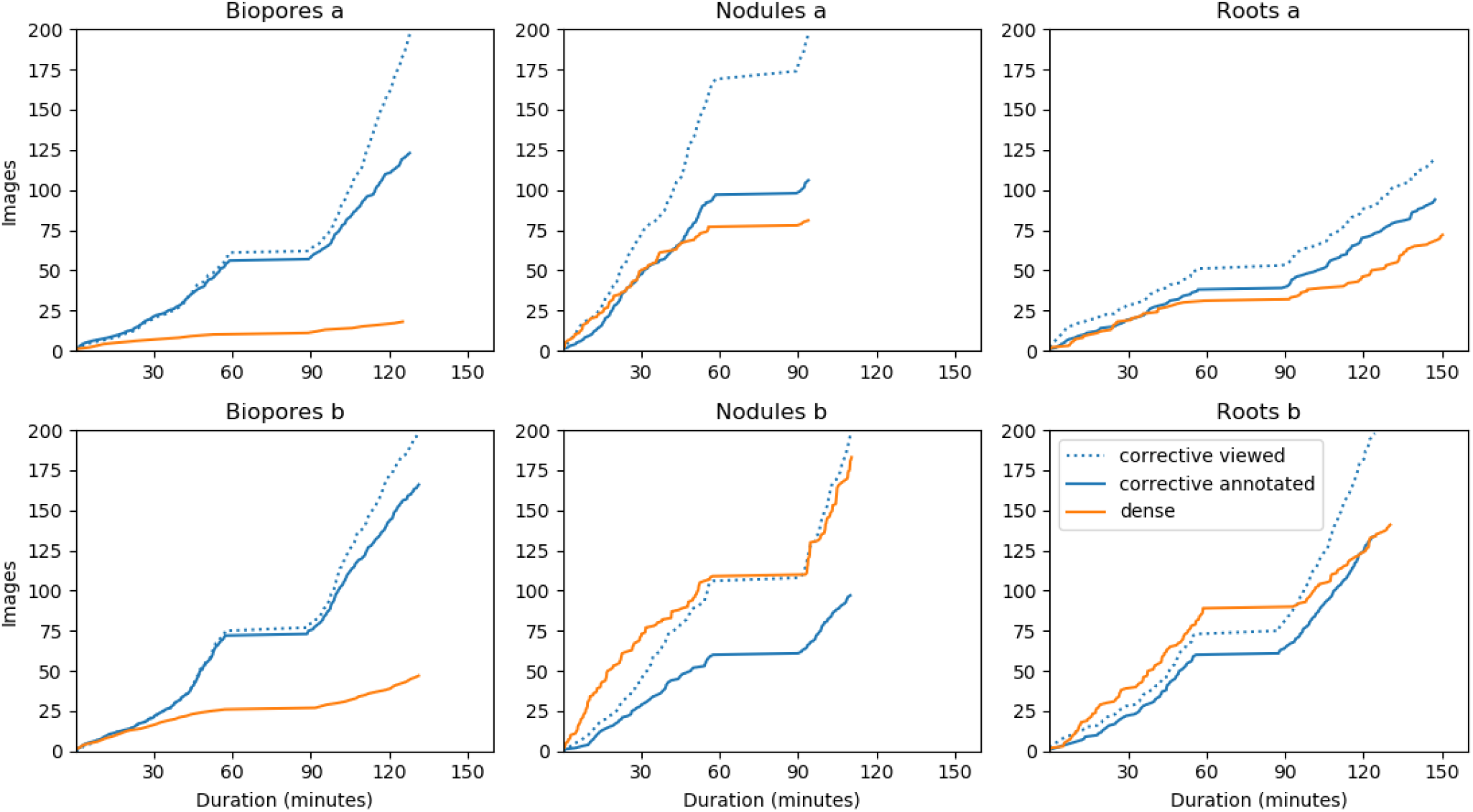
Number of images viewed and annotated for the dense and corrective annotation strategies. For dense all images are both viewed and annotated, where as corrective annotations are only added for images where the model predictions contain clear errors.

For the roots and nodules datasets for user b for the first hour of training, progress through the images was faster when performing dense annotation (Figure 12, Roots b and Nodules b).

We plot the amount of labelled pixels for each training procedure over time for both corrective and dense annotations (Figure 13). With corrective annotation less pixels were labelled in the same time period and as the annotator progressed through the images the rate of label addition decreased (Figure 13).

**Fig. 13.**
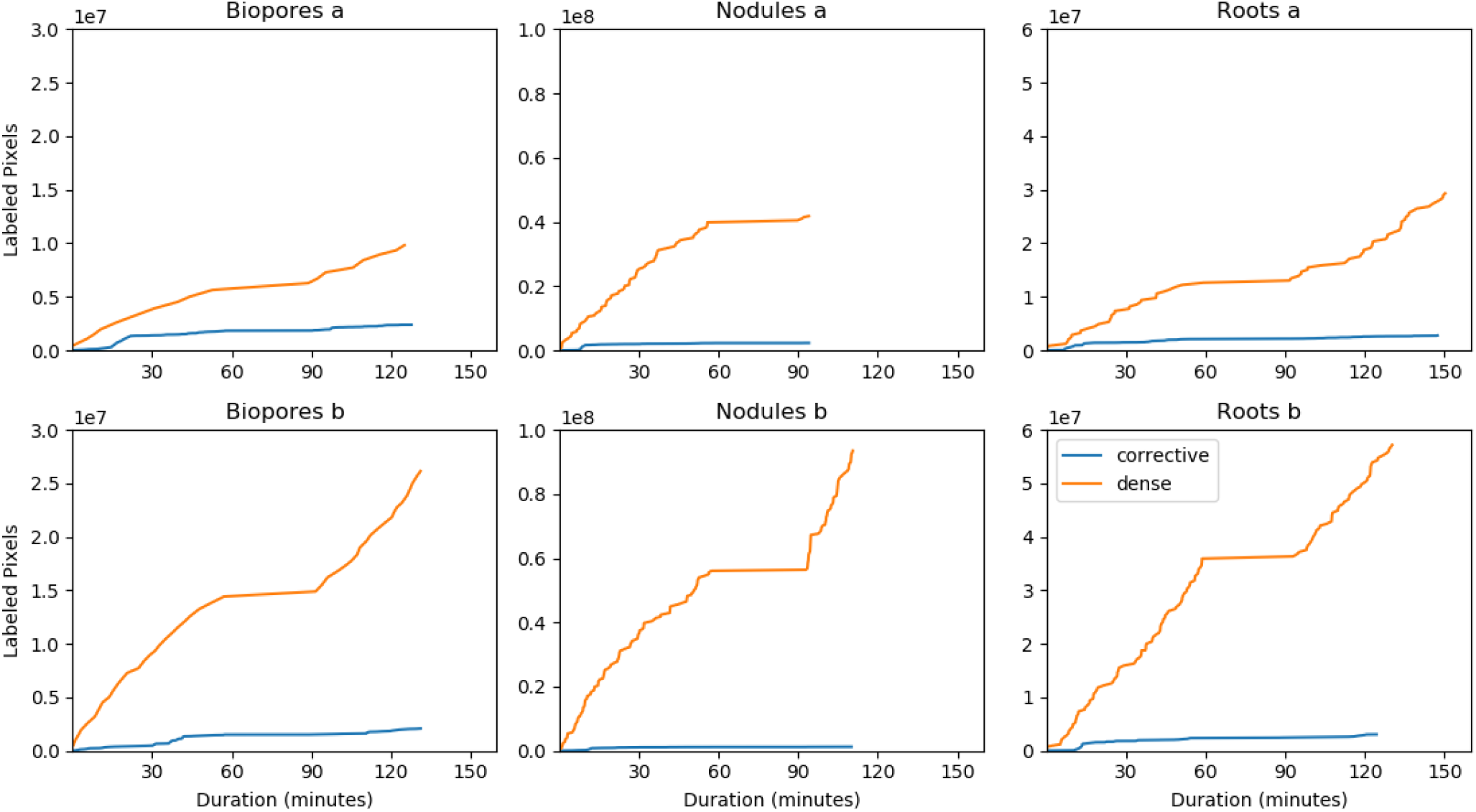
Total number of annotated pixels for dense and corrective annotation strategies over time during the annotation procedure. For dense almost all pixels in each image are annotated. Corrective annotations are only applied to areas of the image where the model being trained exhibits errors.

## Discussion

In this study we focused on annotation duration, as we consider the time requirements for annotation rather than the number of available images to be more relevant to the concerns of the majority of plant research groups looking to use deep learning for image analysis. Our results, for corrective training in particular, confirm our first hypothesis by showing that a deep learning model can be trained to a high accuracy for the three respective datasets of varying target objects, background and image quality in less than two hours of annotation time.

Our results demonstrate the feasibility of training an accurate model using annotations made in a short time period, which challenges the claims that tens of thousands of images (7) or substantial labelled data (69) are required to use CNNs. In practice, we also expect longer annotation periods to provide further improvement. The *R*^2^ for corrective training had a significant correlation with annotation duration indicating that spending more time annotating would continue to improve performance.

There was a trend for an increasing fraction of viewed images to be accepted without further annotation later in the corrective training (Figure 12), indicating fewer of the images required corrections as the model performance improved. This aligns with the reduction in the rate of growth for the total amount of corrections (Figure 13) indicating continuous improvement in the model accuracy over time during the corrective training.

We suspect the cases where dense annotation had a comparatively faster speed in the beginning (Figure 12, Roots b and Nodules b) were due to three factors. Firstly, switching through images has little overhead when using the dense annotation strategy as there is no delay caused by waiting for segmentations to be returned from the server. Secondly, corrective annotation will take a similar amount of time to dense in the beginning as the annotator needs to assign a large amount of corrections for each image. And thirdly, many of the nodule images did not contain nodules meaning dense annotations could be added almost instantly.

Although corrective annotation tended to produce models with higher accuracy relative to dense (Table 3), the lack of a statistically significant difference prevents us from coming to a more substantive conclusion about the benefits of corrective over dense annotation. Despite being unable to confirm our second hypothesis, that corrective annotation provides improved accuracy over dense in a limited time period, it is still clear that it will provide many real-world advantages. The feedback given to the annotator will allow them to better understand the characteristics of the model trained with the annotations applied. They will be able to make a more informed decision about how many images to annotate to train a model to sufficient accuracy for their use case.

Although strong correlation was attained when using the models trained with corrective annotation, they in some cases overestimated (Figure 10 a) or underestimated (Figure 10 b) the objects of interest compared to the manual counts. For the biopores (Figure 10 a) this may be related to the calibration and threshold procedure which results in biopores below a certain diameter being excluded from the dataset. We inspected the outlier in Figure 10 b where RootPainter had overestimated the number of nodules compared to the manual counts. We found that this image (043.jpg) contained many roots which were bunched together more closely than what was typical in the dataset. We suspect this had confused the trained network and could be mitigated by using a consistent and reduced amount of roots per scan, whilst using more of the images for training and annotating for longer to capture more of the variation in the dataset.

In one case, training with corrective annotation failed to produce a model that gave a strong correlation with the manual measurements. This was for the nodules data for user b, where the *R*^2^ was 0.69. We suspect this was partially due to the limited number of nodules in the training data. Many of the images in the dataset created for training contained no nodules and only included the background. This also meant the annotation was able to finish in less time. We consider this a limitation of the experimental design as we expect that a larger dataset which allowed for annotating nodules for the full two hour time period would have provided better insights into the performance of the corrective training procedure.

Figure 6 shows examples of some of the errors in the nodules dataset. In practice, the annotator would be able to view and correct such errors during training until they had abated. We noticed that many of the nodule errors were smaller false positives, so investigated the effect of filtering out nodules less than a certain size (Figure 14). We found correlation increased substantially from 0.69 to 0.75 when changing the threshold from 0 to 5 pixels, which can be explained by the removal of the smaller false positive artefacts (Figure 6).

**Fig. 14.**
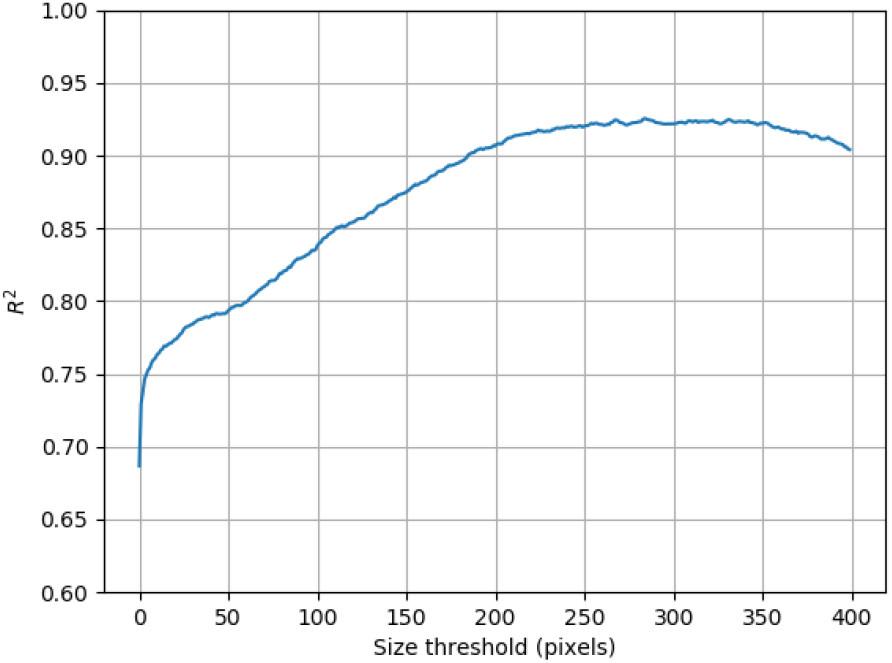
Correlation between automated and manual nodule counting as a function of size threshold for the automatically detected nodules. The thresholded nodules include only those above the specified area in pixels.

The benefits of excluding small nodules continued up to a threshold of 284 pixels, giving an *R*^2^ of 0.93. This indicates that the model was producing many small false positive predictions, which could also explain some of the overestimation of nodules (Figure 10 b).

The problem with small false positives may have been mitigated with the dense annotations as a larger amount of background examples are added, suppressing more of the false positive predictions that arise in the limited training time. The improvement in *R*^2^ when removing small nodules may also be due to differences in subjective interpretation of what is a nodule, between the original counter and annotator training the model.

The reduction in *R*^2^ as dense annotation time increased, shown in nodules b (Figure 11) was highly unexpected. Although in some cases increasing training data can decrease performance when training CNNs (70), it is usually the case that the opposite is observed. We suspect these anomalous results are due to the large amount of variation in the success of the dense training procedure, rather than revealing any general relationship between performance and the amount of data used.

As the nodule images are captured in a controlled environment, further improvements to accuracy could be attained by reducing controllable sources of variation and increasing the technical quality of the images. The lighting was also varying for the nodules with approximately half of the images underexposed. We expect that more consistent lighting conditions would further improve the nodule counting accuracy. Cropping the nodule images manually could also become a time consuming bottleneck, which could be avoided by ensuring all the roots and nodules were positioned inside the border and having the placement of the border be fixed in its position in the scanner such that the cropping could be done by removing a fixed amount from each image, which would be trivial to automate.

Figure 4 indicates corrective annotation leads to lower *R*^2^ in the earlier phases of annotation (e.g. within 60 minutes). We suspect this is due to dense annotation having an advantage at the start as the user is able to annotate more pixels in less time using dense annotation with no overhead caused by waiting for segmentations from the server. We suspect in many cases corrective annotation will provide no benefits in terms of efficiency when the model is in the early stages of training as the user will still have to apply large amounts of annotation to each image, whilst slowed down by the delay in waiting for segmentations. Later in training, e.g. after one hour and 40 minutes, corrective overtakes dense in terms of mean *R*^2^ performance (Figure 4). We suspect this is due to the advantages of corrective annotation increasing as the model converges, when more of the examples are segmented correctly and don’t need adding to the training data as they would provide negligible utility beyond what has already been annotated. Our results show corrective annotation achieves competitive performance with a fraction of the labelled pixels compared to dense (Figure 13). These results align with (71) who confirmed that a large portion of the training data could be discarded without hurting generalization performance. This view is further supported by theoretical work (72) showing in certain cases networks will learn a maximum-margin classifier, with some data points being less relevant to the decision boundary.

The corrective training procedure performance had lower standard error after one hour (Figure 4) and particularly at the end (Table 3). We conjecture that the corrective annotation strategy stabilized convergence and increased the robustness of the training procedure to the changes in dataset with the fixed hyperparameters by allowing the specific parts of the dataset used in training to be added based on the weaknesses that appear in each specific training run.

In more heterogeneous datasets with many anomalies, we suspect corrective annotation to provide more advantages in comparison to dense, as working through many images to find hard examples will capture more useful training data. A potential limitation of the corrective annotation procedure is the suitability of these annotations when used as a validation set for early stopping, as they are less likely to provide a representative sample, compared to a random selection. Our annotation protocol for corrective annotation involved initially focusing on clear examples (Supplementary Note 1) as in preliminary experiments we found corrective annotation did not work effectively at the very start of training. Training start-up was also found to be a challenge for other systems utilising interactive training procedures (27), indicating future work in this area would be beneficial.

Another possible limitation of corrective annotations is that they are based on the model’s weaknesses at a specific point in time. This annotation will likely become less useful as the model drifts away to have different errors from those that were corrected.

One explanation for the consistently strong correlation on the root data compared to biopores and nodules is that the correlation with counts will be more sensitive to small errors than correlation with length. A small pixel-wise difference can make a large impact on the counts. Whereas a pixel erroneously added to the width of a root may have no impact on the length and even pixels added to the end of the root will cause a small difference.

A limitation of the RootPainter software is the hardware requirements for the server. We ran the experiments using two NVIDIA RTX 2080 Ti GPUs connected with NVLink. Purchasing such GPUs may be prohibitively expensive for smaller projects and hosted services such as Paperspace, Amazon Web Services or Google Cloud may be more affordable. Although model training and data processing can be completed using the client user interface, specialist technical knowledge is still required to setup the server component of the system.

In addition to the strong correlations with manual measurements when using corrective annotation, we found the accuracy of the segmentations obtained for biopores, nodules and roots to indicate that the software would be useful for the intended counting and length measurement tasks (Figures 7, 8 and 9).

The performance of RootPainter on the images not used in the training procedure indicate that it would perform well as a fully automatic system on similar data. Our results are a demonstration that for many datasets using RootPainter will will make it possible to complete the labelling, training and data processing within one working day.

## Availability of data and materials

The nodules dataset is available from (66). The biopores dataset is available from (65). The roots dataset is available from (67). The client software installers are available from (73). The source code for both client and server is available from (74). The created training datasets and final trained models are available from (64).

## ACKNOWLEDGEMENTS

Martin Nenov for proof reading and conceptual development. Camilla Ruø Rasmussen for support with using the rhizotron images. Ivan Richter Vogelius and Sune Darkner for support during the final stage of the project. Prof. Dr. Timo Kautz and Prof. Dr. Ulrich Köpke for provision of the biopore dataset. Guanying Chen, Corentin Bonaventure L R Clement and John Kirkegaard for helping test the software by being early adopters in using RootPainter for their root research. Simon Fiil Svane for providing insights into root phenotyping and image analysis.

We thank Villum Foundation (DeepFrontier project, grant number VKR023338) for financially supporting this study.

## Supplementary Note 1: Corrective Training Protocol

### Stage 1

- Start a timer immediately before starting to annotate
- Start training after clicking *Save & Next* for the second annotated image.
- Keep track of how many images you have annotated until you have annotated six images.
- Skip images which do not include clear examples of both classes.
- When images contain clear examples of both classes then label the clear and unambiguous parts of the image.
- Aim to label around 5-10 times as much background as foreground.
- Use a thinner brush to avoid boundaries when labelling the foreground class as these can be ambiguous and time consuming to label.
- After clicking *Save & Next* for the 6th image proceed to stage 2.
- Write down the image number for the 6th annotated image.

### Stage 2

- For each image press S to view the segmentation. Instead of labelling everything which is clear, focus on labeling the parts of the image which have clearly been segmented incorrectly, whilst following the corrective annotation advice.
- Once you have proceeded through 10 images since the 6th annotated image then set pre-segment (from the options menu) from 0 to 1. Increasing the pre-segment setting causes the server to create segmentations ahead of time for upcoming images. This allows the user to progress through the images faster but presents a trade-off as they could potentially be out of date as they are segmented with the best model available at the time and not updated. Thus we only increase pre-segment once the user has worked through a few images, as their annotation time speeds up and necessitates the adjustment.
- Once you have proceeded through 20 images since the 6th annotated image then set pre-segment from 1 to 2.
- Once 1 hour has passed on the timer then take a break for 30 minutes.
- After the 30 minute break then click Start Training again and proceed to annotate as before the break for another hour.
- After the second hour has been completed then stop annotating.
- Leave the network to stop training on its own.

## Supplementary Note 2: Corrective Annotation Advice

- Use a large brush for the background (green) as this makes it quicker label all the false positive regions.
- Focus time and attention on the incorrectly predicted parts of the image
- It is not a problem to label some foreground as foreground which has already been predicted correctly.
- It is also not a problem to label some background as background if it has already been predicted correctly.
- Errors to avoid include labelling a background pixel as foreground or labelling some foreground as background. These should be corrected using the eraser tool.
- It is not a problem to leave small areas unlabelled such as boundaries between foreground and background in the interest of avoiding errors whilst annotating quickly.
- Press I (capital i) to hide and show the image in order to better check the networks segmentation prediction for errors before proceeding to the next image.

## Supplementary Note 3: Dense Annotation Advice

- Set pre-segment (from the options menu) to 10 so that segmentation time does not impact ability to work through the images. Increasing the pre-segment setting causes the server to create segmentations ahead of time for upcoming images. For dense we don’t care about the segmentations so by segmenting 10 in advance it means the client software will never stall their progression through the images because the segmentation has not yet loaded.
- Change the background colour from the default transparency level to a transparency level of 8%. This is because altering the brush transparency allows viewing the object of interest through the background annotation.
- Label each image as all background with a single click using the large brush and proceeded to explicitly annotate all objects of interest (using foreground brush) or ambiguous regions (using the eraser brush) before proceeding to the next image.
- Leave ambiguous regions such as boundaries as undefined, rather than labelling them as foreground or background.
- Once the time limit is reached, use the eraser tool to mark areas not yet annotated in the current image as undefined, stop annotating and click *Start Training*.

## Supplementary Note 4: Server software setup instructions

For our tests we use a client-server architecture and run the client and server components of the system on different computers, using Dropbox to facilitate IO between them. We do not use any Dropbox specific functionality so any service which synchronises a folder between two computers should work. It is also possible to run the client and server on the same computer which will reduce lag and eliminate the need to use a third party service or network drive to sync files. We tested the server component using Python 3.7.5.

1. git clone --branch 0.2.4 https://github.com/Abe404/root_painter.git
2. cd root_painter/trainer
3. pip install torch==1.3.1 -f https://download.pytorch.org/whl/torch_stable.html
4. pip install -r requirements.txt
5. NOTE: pytorch installation may be more involved as it could require and consideration of the current CUDA version. We have tested using pytorch version 1.3.1 but would expect it to work with more recent versions. See https://pytorch.org/get-started/previous-versions/ for more details on how to install pytorch.
6. python3 main.py
7. You will be prompted to specify a sync location. For our tests we used Dropbox and a folder named paper_rp_sync so we input ∼/Dropbox/paper_rp_sync
8. If the folder doesn’t exist then it will be created with the necessary sub folders (datasets, projects, models and instructions).
9. The system will start running and watching for instructions from the client. You must share access to the created folder with the users using your file share service (such as Dropbox) or network drive. The users will then need to input this when they first run the client software.
10. See section *Software Implementation* for further instructions for client software setup.
11. For our tests we ran the RootPainter server inside a tmux session but for more long running use cases a systemd service will likely be more robust. See https://github.com/torfsen/python-systemd-tutorial for instructions on creating a systemd service with python.

## Notes

https://zenodo.org/record/3754074

https://zenodo.org/record/3753603

https://zenodo.org/record/3754081

https://zenodo.org/record/3754046

https://zenodo.org/record/3753969

https://github.com/Abe404/root_painter

### Competing Interest Statement

The authors have declared no competing interest.

### Summary of Updates

Small typos were fixed. The python version is specified for the server. The git tag is also specified with a newer version which has fixes for sshfs file sync.

